# Identifying drug sensitivity of multifocal primary prostate cancer

**DOI:** 10.1101/2023.12.12.569525

**Authors:** Juening Kang, Panagiotis Chouvardas, Katja Ovchinnikova, George N. Thalmann, Sofia Karkampouna, Marianna Kruithof-de Julio

## Abstract

The pronounced intra-patient variability in multifocal primary prostate cancer (PCa) has curtailed the efficacy of current treatment options. Patient-derived organoids (PDOs) have emerged as a pivotal model for functional testing due to their capability to retain the histopathological and molecular characteristics of parental tissues, allowing timely acquisition of drug response outcomes. In our study, employing twin biopsies from multiple lesions with matched PDO models *in vitro*, we investigated the molecular heterogeneity of PCa, and how it is linked to in vitro PDO pharmacological heterogeneity. Our functional testing approach leverages PDOs to screen standard-of-care treatment and FDA-approved compounds for other malignancies, aiming to repurpose their use in PCa and explore alternatives to androgen deprivation therapy. By integrating gene expression data from parental tissue with drug response results from PDOs, we have established a transcriptomics-based drug prediction models. The machine learning-based prediction model can predict the experimental PDO response to a specific drug, for the majority of screened drugs. This study offers a preclinical approach to potentially procure drug prediction outcomes and validate them in PDO models, as a prior step to clinical trial investigations or for selection of targeted therapeutic options.

## Introduction

Prostate cancer (PCa) is the second most common cancer in men worldwide, with manifestations varying from indolent localized tumours to widespread metastases, indicating a highly heterogeneous nature^1-3^. At the time of diagnosis, 60-90% of PCa patients are found to have multiple lesions with different Gleason scores (GS), DNA ploidy, and aggressive phenotypes, which can develop independently in the same prostate and pose a therapeutic challenge^4,5^. The current standard treatments for PCa are limited in terms of long-term effectiveness due to targeting the cancer as a bulk, without considering the multifocal nature of PCa and the intrinsic heterogeneity of each lesion^6^. Therefore, understanding the molecular (transcriptomic and genomic) heterogeneity of PCa and its impact on treatment response first at the cellular level and subsequently at the patient level is critical for optimizing personalized precision medicine decision.

Recent studies have revealed that somatic mutations or DNA copy number changes are rarely shared among different tumour lesions within the same prostate^7^, but also within a given lesion^8^. The presence of high inter- and intra-tumoural genomic heterogeneity and the sub- clonal diversity influences gene expression among distinct foci^9,10^. Current tools fail to identify which properties of tumour foci are associated with higher risk for metastatic progression and /or lethal PCa. Such molecular complexity may affect treatment efficacy to androgen deprivation therapy (ADT)^11^. For instance, the aberrant isoforms of androgen receptor AR (AR- v7) in different lesions may lead to evasion from the therapeutic effects of anti-AR therapies, which are generally effective, leading to resistance to second generation AR inhibitors^12,13^. In fact, it has been shown that greater genomic diversity at the primary stage^14^ is associated with resistance to ADT in patients. Molecular evolution analyses of independent tumour foci showed differential drug sensitivity to neoadjuvant ADT among the different foci in case report studies^15^. Inclusion of larger cohorts to dissect the responses of specific lesions pre- and post- treatment would be highly informative.

Such exploration of direct drug-mediated biologic responses in PCa, has been hampered by the slow progression of the disease and lack of suitable cell models, depicted by the limited number of PCa cell lines available, especially for primary stage, AR+ and treatment-naive PCa^16,17^, with the exception of 22Rv1 or E006AA^18,19^. This limitation can be circumvented using patient-derived organoids (PDOs), which have the potential to recapitulate cell type composition and tumour-specific molecular features. Yet they require fresh tissue which is incompatible with routine histopathologic analysis that is prioritized for diagnostics. Previously, we, among others, have shown that PDOs from primary PCa may serve as cellular model similar to primary cultures, which although not indefinitely proliferative, show higher establishment efficiency compared to i.e. 2D cell lines, and thus suitability for preclinical short-term patient-oriented studies^20-23^.

Given the early metastatic nature of PCa (70% with bone metastases)^24^, identification of primary PCa traits, such as molecular features and therapeutic vulnerabilities is crucial for patient stratification and therapeutic management. In this study, a precision medicine approach of investigating functional drug responses of individual foci, in parallel with genomic and gene expression in twin biopsies, has been employed. Using our previously developed methodology for PDO derivation from primary and advanced PCa to assess *in vitro* chemosensitivity to standard-of-care (enzalutamide) and to a selected drug candidate panel of non-ADT related compounds that showed efficacy in PCa^21^. Multifocal features of PCa, including histological and molecular heterogeneity, as well as non-malignant epithelial areas were analysed to obtain information on subtle molecular possibly premalignant features. To further explore predictors of drug response, we analysed the association between drug sensitivity or resistance and different parameters such as genomic mutations, gene expression, and clinical parameters. Machine learning (ML) models were built and trained to predict drug response in multifocal PCa patients, aiming to provide additional insights for personalised precision medicine treatment decision making.

## Results

### Generation and culture of PDOs from heterogenous primary PCa clinical samples

Tissues from radical prostatectomies were divided in quadrants and four core biopsies were obtained from each quadrant. The cores were obtained as twin/mirror biopsies; one half of the core was processed for histology and omics, and the other half was processed for cell derivation. Our entire cohort consists of 24 patient cases with at least one tumour core (GS≥6). PDOs were generated from four distinct areas (designated as cores A-D) of each prostate (**Fig. 1A**). To investigate the histological heterogeneity, we compared the histopathological features of the distinct foci at the tissue level. As shown in **Fig. 1B**, diverse histopathological features exhibited in distinct foci. The majority of patient tissues (83.3%, N=20/24) had a GS of 7 and a 12.5% (N=3/24) of GS 8-10 (**Sup. Fig. 1A**, *left*). Varying GS and tumour content emerged when assessing distinct foci from each patient with 28.1% of GS7 (N=27/96), and 14.6% in GS8-10 (N=14/96) (**Fig. 1C, Sup. Fig. 1A**, *right*). This indicated that the GSs determined based on a single lesion might not comprehensively depict the patient’s disease stage.

**Figure 1.**
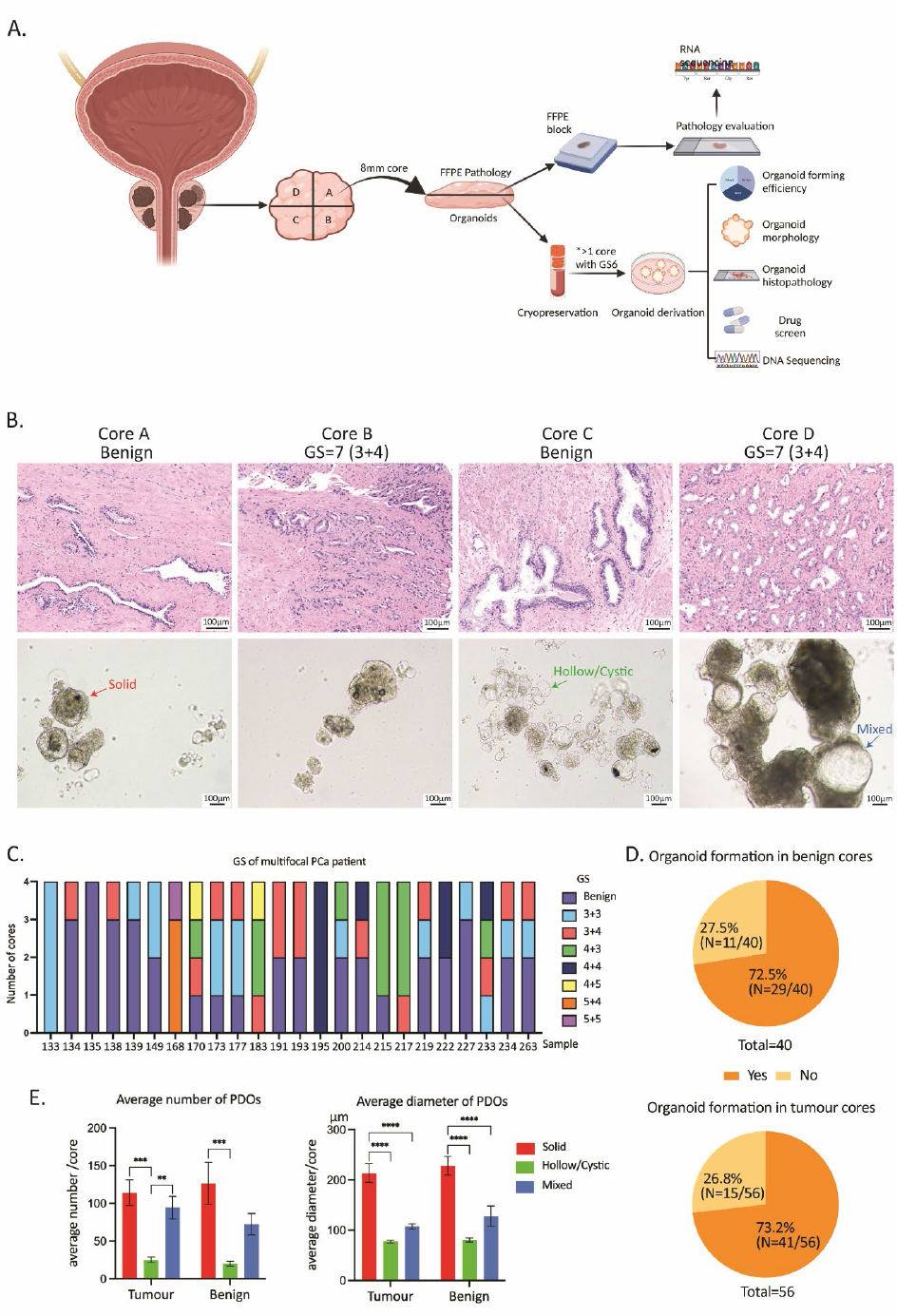
Derivation and culture of patient-derived organoids (PDOs) from multifocal primary prostate cancer (PCa) (A) Scheme of the experimental protocol for multifocal primary PCa organoid derivation and analysis. (B) Morphology of multifocal primary PCa and matched PDOs (passage (p) 0) for one representative case. H&E images of primary tumour and brightfield images of deriving PDOs. (C) Gleason score (GS) reported to each core of the patient. (D) Ratio of PDO formation and no PDO formation over the total samples cultured for histopathologically defined benign cores (n=40) and histopathologically defined tumour cores (n=56) (unpaired t test, *p*=0.2658). (E) Quantification (average number and diameter/core) of PDO morphology on histopathologically defined benign or tumour cores (average number of solid morphology vs. hollow morphology in tumour-containing cores: *p*=0.0002; average number of solid morphology vs. hollow morphology in non-tumour-containing cores: *p*=0.0002; average diameter of solid morphology vs. hollow and mixed morphologies in tumour-containing cores: *p*<0.0001; average diameter of solid morphology vs. hollow and mixed morphologies in non-tumour-containing cores: *p*<0.0001).

PDO forming efficiency in relation to parental tumour tissue was investigated. PDO cultures were successfully established from PCa samples irrespective of tumour content, with no significant differences in organoid-forming efficiency between non-tumour-containing cores (N=29/40, 72.5%) and tumour-containing cores (N=41/56, 73.2%, two-sided Fisher’s test, *p*-value > 0.99, **Fig. 1D**). PCa PDOs formed within 5-11 days, while failure to generate PDOs did not correlate with initial number of viable cells (**Sup. Fig. 1B**, unpaired t test, *p*-value=0.2658). PDOs were characterized by morphological analyses and grouped into three morphological patterns: solid, hollow, and mixed. Solid organoids were densely packed epithelial structures without a lumen, hollow organoids consisted of a lumen surrounded by a layer of cells, while mixed structures presented both solid and hollow features (**Fig. 1B**). Despite PDOs generated from different foci exhibited varying morphological characteristics, solid structures were predominantly presented in distinct foci of the same patient, irrespective with parental tissue histopathological features (**Fig. 1B** and **E**). We explored the correlation between different PDO morphologies and histopathologic features of the matched tissue. We calculated the average number and diameter of each PDO morphology that presented from either tumour or benign cores. Solid PDOs were the most prevalent morphology and of larger diameter, irrespective of tissue origin (tumour/benign) (**Fig.1E**). Tumour-containing cores with high GS and tumour content showed a trend for higher number of PDOs with solid and mixed features, while PDOs derived from cores with GS of 7 showed significantly larger diameter compared to tissues with GS of 6 and 9 (two-way ANOVA, *p*-value<0.0001, *p*-value=0.0084 respectively) (**Sup. Fig. 1C-D**).

### PDOs retain histological features of parental tissues

To compare the histological features of PDOs with their parental tissue, as well as between tumour and non-tumour containing cores, the expression of prostate epithelial lineage markers in both PDOs and their parental tissue were assessed through immunofluorescence (IF) staining. We investigated an array of markers, encompassing epithelial (E-Cadherin), luminal (CK8 and AR), basal (p63 and CK5), and the proliferation marker (Ki67) (**Fig. 2**).

**Figure 2.**
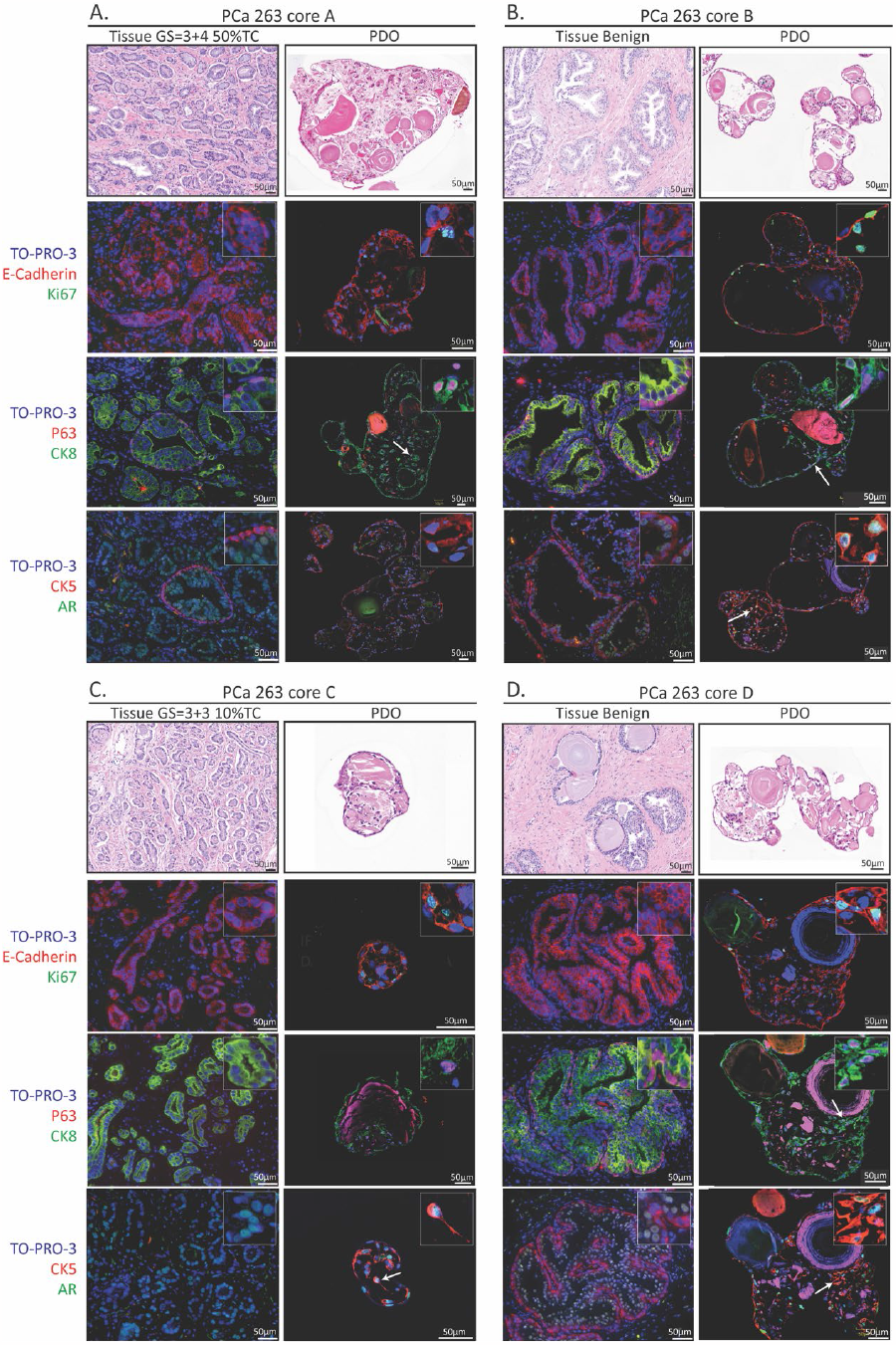
Prostate cancer (PCa) patient-derived organoids (PDOs) originating from independent tumour/benign foci recapitulate histopathological features *in vitro*. H&E and immunofluorescent staining of PDOs and parental tissue for indicated markers. Four representative cores histopathologically defined tumour (A and C, PCa263 core A and core C) and histopathologically defined benign (B and D, PCa263 core B and core D). White arrows indicate examples of cells co-expressing both basal and luminal markers.

The expression of E-Cadherin marker vouched for the epithelial origin of the prostate gland in both the parental tissue and PDOs. The presence of Ki67-positive cells in PDOs indicated their proliferative capacity; however, their absence in parental tissue might stem from the low proliferative index of primary PCa compatible with the slow disease progression.

The expression of both luminal (CK8 and AR) and basal (P63 and CK5) markers substantiated the co-existence of the two prostate epithelial lineages in the organoid cultures. Furthermore, luminal CK8 and AR positive cells were identified around a lumen-like gland in PDOs, surrounded by basal P63 and CK5 positive cells. This was consistent across benign tissues, as well as PDOs from both tumour and benign tissues, although tumour tissues exhibited predominant luminal (CK8 and AR positive) cells with a few surrounding basal (P63 and CK5 positive) cells (**Fig. 2A, C**). Intriguingly, an intermediate cell population, referred to as intermediate or transit amplifying cells^25,26^, that co-expresses luminal and basal markers (CK5 and P63, AR and CK5) was also identified in organoids (**Fig. 2**, *white arrows*), indicating an intermediate differentiation potential.

### Heterogeneous PDO drug responses may be predicted using a transcriptomics-based ML model

Considering the intrinsic heterogeneous histological and molecular features observed in primary PCa, different tumour foci within the same prostate may possess distinct biological properties and molecular features, which may consequently result in diverse response to therapeutic drugs. Therefore, we sought to determine if there is also functional variability in terms of drug response. To address this, we conducted drug screening on PDOs using 11 selected compounds based on our previous study^21^, including standard-of-care (SOC, Enzalutamide) as well as different FDA-approved drugs with indications for other solid cancers such as tyrosine kinase Inhibitors (TKIs, Bosutinib, Crizotinib, Ponatinib, etc.), Anthracyclines (Daunorubicin, Doxorubicin, and Epirubicin), and mTOR inhibitors (Rapamycin, Temsirolimus). These compounds previously showed efficacy in organoids from primary and advanced patient samples and patient-derived-xenografts of PCa. In this cohort, three TKIs (Crizotinib, Ponatinib, and Sunitinib) demonstrated broad effectiveness representing the most effective drugs. Conversely, Enzalutamide, along with mTOR inhibitors (Rapamycin and Temsirolimus) and EGFR inhibitor Erlotinib, indicated an overall lower efficacy in reducing PDOs viability (**Fig. 3A**).

**Figure 3.**
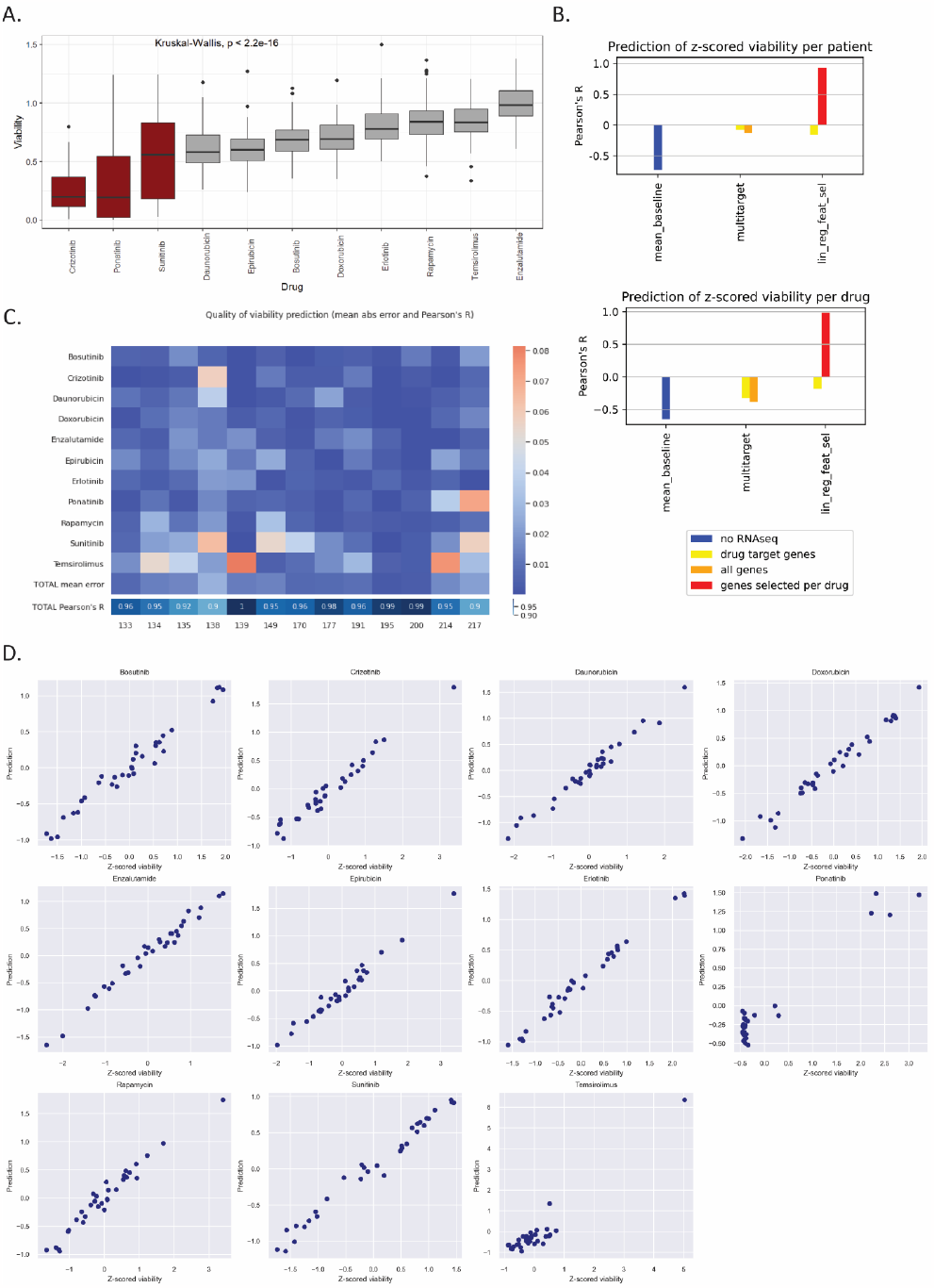
Drug response prediction in multifocal primary prostate cancer (PCa) (A) Results of drug screen assay from 63 PDOs indicate high efficacy of Receptor Tyrosine Kinase Inhibitors (Crizotinib, Ponatinib, Sunitinib) in reducing organoid viability. The ATP-mediated cell viability values were distributed across our cohort (n = 63 PDOs from distinct tumour/benign cores derived from 24 different patient samples) and the drug panel tested. The most effective compounds in the majority of the cases tested, are highlighted in red. Statistical test Kruskal-Wallis. (B) Mean Pearson’s R correlation between predicted and observed drug response values for the models predicted for each drug (left) and each patient (right). The colours of the bars show the RNAseq features used in each model. (C) Mean prediction error of liner regression model with selected features for drugs/patients. The heatmap above shows the mean prediction error for each patient’s response to each drug. The heatmap below shows the Pearson’s R correlation between predicted and observed drug response values for each patient. (D) Pearson’s correlation between predicted and observed drug response values for each drug. The dot plots report the predicted (y axis) and observed (z-scored viability, x axis) values for each core’s response to each drug.

We investigated the presence of transcriptomic diversity of multifocal primary PCa, by subjecting the parental tissues of tumour foci and benign areas to RNA-sequencing (RNA-Seq). To improve the output of RNA-Seq impacted by the RNA degradation in the FFPE tissues, we used random primer cDNA synthesis in combination with targeted sequencing by hybridization with 5’ exon-specific probes. Higher number of mapped reads and exonic regions was achieved with this method, compared to ribosomal RNA depletion and total RNA-Seq. We next explored whether the transcriptomic profile of the tumour tissue (cores) can be indicative of the in vitro drug response in PDOs derived from the matched cores. We sought to develop a machine learning model for predicting overall drug response, but also to individual drugs specifically. To achieve this, we employed several drug-specific standard models and trained them using a “leave one out” approach, leveraging data on both original cell viability and z-scored cell viability for each drug response.

The linear regression model with selected features using Pearson’s correlation relevance score demonstrated standout performance in z-scored cell viability settings (**Fig. 3B**). On average, approximately 260 genes per drug were chosen for the viability setting, while around 1800 were selected for the z-scored viability setting. Intriguingly, we discerned a fluctuation in prediction quality. It peaked when several hundred genes were included as features and then waned as further gene features were integrated. Consequently, we pinpointed the optimal number of genes—where prediction quality was at its zenith—for each drug and employed them as the feature set to train the ML model. Specifically, for Temsirolimus, a high Pearson’s R value (>0.8) was achieved using only a minimal set of features. This high correlation can be traced back to a distinct outlier that was discernible with a few genes, as depicted in **Fig. 3D**, precise prediction of this outlier significantly bolstered the overall Pearson’s R value. Inclusion of other features such as preoperative clinical and histopathological parameters did not further improve the predictions, highlighting that the transcriptomic profile is a sufficiently defining factor.

To assess the predictive accuracy per patient and drug, we undertook Pearson’s correlation analyses, comparing actual observed cell viability values to the model’s predictions. The prediction errors between predicted and actual values consistently hovered below 0.08 for all patients and drugs (**Fig. 3C**). Strong Pearson’s correlations were evident between the model’s predictions and the actual outcomes for every patient (**Fig. 3C**). This suggests our model’s high accuracy in foretelling drug responses on an individual basis. Moreover, scatter plots revealed a pronounced linear relationship between predicted and observed viability values, except for the outlier in the Temsirolimus data (**Fig. 3D**). While expanding the sample size would further fortify the reliability and robustness of our predictions, our pioneering ML model underscores the potential of ML to revolutionize precision oncology and clinical decision-making.

## Discussion

In this study, we elucidated the molecular and functional heterogeneity of primary PCa and the potential mechanisms underlying drug response, utilizing PDOs. Additionally, we developed a transcriptomics-based machine learning drug prediction models, offering a preclinical approach with the potential to inform and enhance clinical trials aimed at targeted therapeutic investigations.

Overall, our patient cohort is well representative of PCa heterogeneity, encompassing different clinical stages with multifocal lesions ranging from GS 6 to GS 10. Although there was a consistent rate of PDO formation and a predominant solid PDO morphology observed across tissues with varying histopathological features, the inherent histological and genetic heterogeneity of PCa were still retained in PDOs. This retention is underscored by the expression of both luminal and basal markers in PDOs, aligning with findings that suggest both cell types may be precursors to varying PCa stages^27-30^. The tumour heterogeneity within PCa necessitates a complex molecular classification of patients^31,32^.

Gene expression profiling of independent tumour foci can bridge the gap between genomics and phenotype and identifying molecular features associated with tumour or benign states. PDX-derived organoids and PDOs show high trancriptomic correlation with their originating tumour tissues as we previously have shown^21^. Thus, we hypothesized that RNA sequencing on the parental tissue in combination with PDO drug testing may represent a suitable approach to evaluate drug efficacy and to prioritize compounds for in vitro validation, but also to associate gene expression-drug associations and identify drug vulnerability mechanisms. From our previous study, a PDX-organoid automated screen revealed efficacy of several TKI drug compounds (e.g. Bosutinib, Sunitinib, Ponatinib) in primary and advanced PCa^21^. In the present study, we highlighted the consistent high efficacy of Crizotinib, which stood out among 3 most effective TKIs (Crizotinib, Ponatinib, and Sunitinib) observed in our cohort across heterogenous tumour lesions. Phosphoproteomics have revealed a higher tyrosine kinase phosphorylation activity in primary PCa versus benign tissues, as well as distinct phosphorylation signatures that are stage-specific “footprints” for primary or CRPC stage ^33,34^, which explains the efficacy of different TKIs in our PDO cohort targeting multiple Receptor Tyrosine Kinases. TKIs are increasingly being considered for and shown efficacy in primary PCa treatment^33,35^.

As a target for Crizotinib, c-Met is not only confirmed to be overexpressed in primary PCa^36^, but also exhibits an inverse correlation with AR expression^37,38^. Given that ADT remains the SOC for PCa, the removal of androgens in PCa patients could further augment c-Met expression and driving cancer progression^39^. This further supports the effectiveness of Crizotinib in PCa cell lines, organoids, and PDX models^21,40,41^. Although the efficacy of Crizotinib is mostly observed in the treatment of advanced PCa, our PDO post-treatment RNASeq indicated potential mechanisms by which Crizotinib inhibits primary PCa cell growth. This provides novel insights into the potential application of Crizotinib for PCa treatment^42,43^. Despite the general efficacy exhibited by the three TKIs in our cohort, the spatial molecular variations and varied drug sensitivities in primary PCa emphasize the importance of matching spatial sampling with drug response^14,15^. Furthermore, there is a clear need to evaluate drug responses from multiple lesions comprehensively, as these insights will be instrumental in guiding treatment decision-making. Given the timeliness and capability of PDOs to capture the characteristics of parental tumours, we have, in conjunction with drug response data from PDOs and transcriptomic profiles from parental tissue, established a transcriptomics-based ML model. Various combinations including histopathological features, genomic profiles, and clinical parameters were explored during the algorithm development. However, only transcriptomics consistently emerged as the most effective contributor to predictions^44,45^. This might be attributed to the capacity of transcriptomic features to more comprehensively characterize intrinsic tumour signalling patterns, enabling the ML model to better capture drug target information^46,47^. The transcriptomics-based ML model, using RNASeq data from FFPE specimens, allow us to predict the collective response across various drugs for multifocal lesions and the response of individual tumour and benign foci to individual drugs and to evaluate drug response heterogeneity. Using RNA-Seq data obtained on FFPE specimens, is compatible with routine diagnostic practises and subsequently a more targeted panel would be likely to be developed.

The present study aimed to has deepen the understanding of the heterogeneity in multifocal PCa, particularly its functional variability. While PDOs have shown promise for preclinical drug validation, most of the drugs we tested have not been FDA-approved for PCa treatment. As such, our drug response and prediction findings cannot be directly applied to patients at present. However, this limitation may soon be alleviated with ongoing Phase I clinical trials for Crizotinib in advanced PCa (ClinicalTrials.gov #NCT02207504), in combination with enzalutamide (NCT02465060), for Ponatinib and other drugs included in our panel for CRPC treatment (Erlotinib, Doxorubicin, Everolimus, #NCT03878524), as well as the promising results Sunitinib has already achieved in clinical trials (#NCT00631527 and # NCT00672594). In summary, this study provides a detailed exploration of the complex molecular and functional aspects of multifocal primary PCa. It illuminates the correlation between gene expressions and drug response, offering potential biomarkers and therapeutic targets for future investigations. Moreover, it introduces ML models that assist in clinical decision-making, significantly advancing the evolving landscape of PCa treatment and contributing to the progress of precision medicine.

## Acknowledgements

The authors would like to thank the Institute of Tissue Pathology in Bern, the study nurse and medical personnel at the Department of Urology, Inselspital Bern, and the patients who participated in this study.

