## Supplementary Figure 1 for "Identifying drug sensitivity of multifocal primary prostate cancer"

### Supplementary Material

#### Supplementary Figure 1

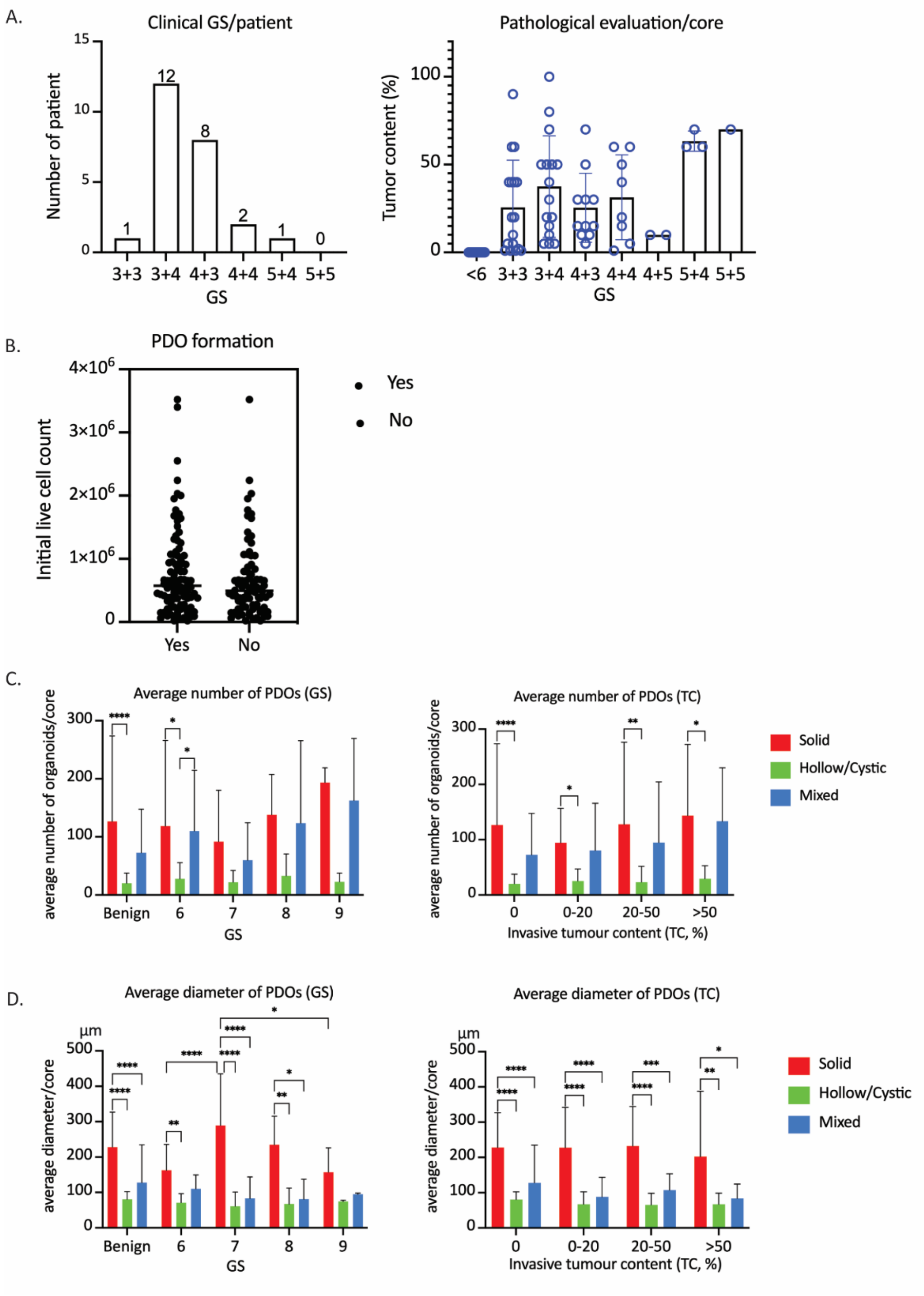


**Sup.Fig 1. Histopathological characteristics of tissue, and PDO quantification and forming efficiency.**

(A) Overall clinical Gleason score (GS) reported per patient (left) and pathological evaluation reported reported per core (right).

(B) PDO formation or no formation regarding initial live cell count per core (unpaired t-test, p-value=0.2658)

(C-D) Quantification (average number and diameter/core) of PDO morphology on histopathologically defined benign or tumor cores, with varying GS and tumour content (*p-value<0.5, **p-value<0.1, ***p-value<0.01, ****p-value<0.001).
